# New insights into the *cis*-regulation of the *CFTR* gene in pancreatic cells

**DOI:** 10.1101/2025.01.14.632890

**Authors:** Clara Blotas, Anaïs Le Nabec, Mégane Collobert, Mattijs Bulcaen, Marianne S. Carlon, Claude Férec, Stéphanie Moisan

## Abstract

**Background:** Spatial organization of the genome is fundamental for ensuring accurate gene expression. This process depends on the communication between gene promoters and distal *cis*-regulatory elements (CREs), which together make up 8% of the human genome and are supported by the chromatin structure. It is estimated that over 90% of disease-associated variants are located in the non-coding region of the genome and may affect CRE. For the *cystic fibrosis transmembrane conductance regulator* (*CFTR)* gene, a complete understanding of tissue-specific *CFTR* expression and regulation is missing, in particular in the pancreas. Mechanistic insights into tissue-specific expression may provide clarity on the clinical heterogeneity observed in Cystic Fibrosis and CFTR-related disorders.

**Methods:** To understand the role of 3D chromatin architecture in establishing tissue-specific expression of the *CFTR* gene, we mapped chromatin interactions via circular chromosome conformation capture (4C) and epigenomic regulation through H3K27ac and DNase Hypersensitive site I (DHS) in Capan-1 pancreatic cells. Candidate regulatory regions are validated by luciferase reporter assay and CRISPR-knock out.

**Results:** We identified active regulatory regions not only around the *CFTR* gene but also outside the topologically associating domain (TAD). By performing functional assays, we validated our targets and revealed a cooperative effect of the −44 kb, −35 kb, +15.6 kb and 37.7 kb regions, which share common predicted transcription factor (TF) motifs. Comparative 3D genomic analysis and functional assays using the Caco-2 intestinal cell line revealed the presence of tissue-specific CREs.

**Conclusion:** By studying the chromatin architecture of the *CFTR* locus in Capan-1 cells, we demonstrated the involvement of multiple CREs upstream and downstream of the *CFTR* gene. We also extend our analysis to compare intestinal and pancreatic cells and provide information on the tissue-specificity of CRE. These findings highlight the importance of expanding the search for causative variants beyond the gene coding sequence but also by considering the tissue-specific 3D genome.

## Background

Over the last decades, functional genomics has become increasingly important, particularly with the development of new technologies. They allows to gain insights about the unexplored part of the genome, *i.e.* the non-coding genome (1). The ENCODE (Encyclopedia of DNA Elements) project, launched in 2003, aims to provide a comprehensive annotation of the entire genome. The consortium estimated that 8% of the genome consists of candidate *cis*-regulatory elements, known as cCREs (2). CREs are regions bound by transcription factors (TFs) that drive the cell type-specific expression of target genes regardless of their orientation and genomic distance. They can be classified as enhancers, silencers, insulators or promoters, active or poised. CREs are characterized by biochemical marks such as open chromatin, nucleosome-free regions and post-translational modifications (PTMs), such as acetylated histone H3 Lys27 (H3K27ac) for active enhancers, trimethylated histone H3 Lys4 (H3K4me3) for promoters, and CCCTC-binding factor (CTCF) binding for insulators (3). To be functional, the regulatory region must interact with its target gene.

*Cis*-regulation is a highly conserved mechanism that is essential for correct spatio-temporal gene expression (4). Tissue and temporal specificity make *cis*-regulation highly dynamic and challenging to study. Further research is however needed, both to improve knowledge and to apply these new insights to human diseases. Most variants causing genetic diseases are thus far located in protein coding regions, but whole-genome studies are providing increasing evidence that variants in non-coding regions are involved in pathogenesis or are at risk in specific tissues (5,6). It is estimated that over 90% of disease-associated variants are located in the non-coding region of the genome and may have an impact on CRE (7).

CFTR-associated diseases include cystic fibrosis (CF) and CFTR-related diseases (CFTR-RD) (8). These conditions result from variations in the *CFTR* gene, which lead to aberrant expression or dysfunction of the encoded protein. The *CFTR* gene encodes CFTR, an ion channel responsible for conductance of chloride and bicarbonate that is expressed at the apical membrane of epithelial cell layers in several organs. Its absence or dysfunction causes a loss of ion homeostasis at the epithelial cell surface, resulting in mucus disturbance, organ obstruction, and chronic bacterial infection. Without appropriate care, these conditions can lead to death during infancy.

Significant genotypic and phenotypic heterogeneity is observed in CFTR-associated diseases, with over 2100 variants and a spectrum of clinical presentations ranging from mild to severe. CF affects multiple organs, including the lungs, small intestine, pancreas, reproductive tract, and liver. In contrast, CFTR-RD are clinical entities with features of CF and evidence of CFTR dysfunction but do not meet the criteria for a CF diagnosis (9). The majority of these diseases manifest as mono-organ forms, including CBAVD (congenital bilateral absence of vas deferens), pancreatitis, and bronchiectasis (10). The complexity of the clinical picture cannot be fully explained by variants in the coding sequence alone, and the regulation of the *CFTR* gene remains incompletely understood. Insight into the *cis*-regulation of the *CFTR* gene may explain at least part of this phenotypic variability.

The *CFTR* gene is surrounded by two CTCF TAD boundaries located at −80.1 kilobases (kb) upstream of the transcription start site (TSS) and +48.9 kb downstream of the last *CFTR* codon. The first description of *CFTR* regulatory elements occurred in 1996 with the identification of a DNase I hypersensitive site (DHS) bound by TFs with enhancer activity within intron 1 of the *CFTR* gene (11). Subsequent studies have focused on the identification of pulmonary and intestinal CREs on the basis of the severity of symptoms in these tissues and the availability of models. A comprehensive review of previous studies is provided in (12). With respect to the pancreas, although there is a lack of knowledge and relevant models, more studies are needed, as pancreatic insufficiency represents a major manifestation in 85% of people with CF (pwCF). Additionally, pancreatitis is the second most common CFTR-related disorders (CFTR-RD).

With this in mind, in this work, we combine chromatin study techniques, including 4C-seq, ATAC-seq and CUT&RUN-seq (Cleavage Under Targets & Release Using Nuclease), to identify putative CREs in the pancreas, where little information is currently available, as recommended by Gasperini *et al.*(3). The activity of the regulatory regions was assessed via a luciferase reporter assay and we have started validating our target regulatory regions through CRISPR knock-out experiments. We identified regulatory regions previously shown to be involved in other tissues, as well as newly described regions, both within and outside the TAD. Overall, this work provides a more comprehensive regulatory landscape of the *CFTR* gene in the pancreas and highlights the tissue-specific regulation of this gene.

## Methods

### Cell Culture

Three cell lines were used: Capan-1, which is derived from pancreatic adenocarcinoma, Caco-2, which is derived from colorectal adenocarcinoma and HepG2, which is derived from hepatocellular carcinoma. Cells were grown in Dulbecco’s modified Eagle medium supplemented with 10% fetal bovine serum. The samples were incubated at 37°C with 5% CO_2_. The cells are frequently tested for mycoplasma contamination.

### ATAC-seq

Omni-ATAC-seq libraries were generated as previously described (13). A total of 5.10^4^ pelleted cells were resuspended in cold lysis buffer and incubated on ice for 3 minutes. Transposition was performed with buffer containing tagment DNA enzyme (Illumina) at 37°C and 1000 rpm for 30 minutes. DNA was purified via a MinElute PCR purification kit (Qiagen). Libraries were amplified via NEBNext High-Fidelity 2 × PCR Master Mix (New England Biolabs) and Nextera adapters (Illumina). The PCR products were purified via a QIAquick PCR purification kit (Qiagen). Libraries were quantified via Qubit (Thermo Fisher), and fragment sizes were analyzed via a bioanalyzer (Agilent). Libraries were paired-end sequenced via Illumina sequencing (NextSeq500, Illumina). The generated paired FastQ reads were analyzed via the atacseq pipeline version 0.12 with default parameters and mapped to the hg19 reference genome (https://github.com/iwc-workflows/atacseq). After filtering and duplicate removal, peaks are called via macs2. A BigWig file contains the coverage file. For HepG2 cells, we downloaded ATAC-seq fastQ from (14) (GSE 139190).

### CUT&RUN-seq

CUT&RUN-seq libraries were generated as previously described (15). A total of 5.10^5^ cells were incubated with activated concanavalin A beads (Bangs Laboratories) for 10 minutes at 4°C. The cells were resuspended in buffer containing 0.06% digitonin (Calbiochem), and specific antibodies against H3K27ac (Ab4729), CTCF (Active Motif 61311), and H3K27me3 (Millipore 07-449) were added and incubated at 4°C for 2 hours. pAG-MNase (1 ng/µL, Cell Signaling Technology) was added, and the mixture was incubated for 1 hour on ice. Unbound enzymes were washed, and calcium chloride (2 mM) was used to activate pAG-MNase for 30 minutes on ice. After incubation, stop buffer was added, and the samples were incubated at 37°C for 30 minutes to release DNA fragments. Purification was performed via DNA purification buffers and spin columns (Cell Signaling Technology). End-repair and A-tailing were then performed via the KAPA HyperPrep Kit (Roche). Next, 10 µL of 4X ERA buffer were added to the libraries, which were subsequently incubated at 20°C for 30 minutes and at 50°C for 60 minutes. Ligation of KAPA UDI adapters (0.3 µM) was performed at 20°C for 1 hour. Size selection was performed via AMPure XP beads (Beckman Coulter) with double purification via the addition of HXP buffer (20% PEG 8000, 2.5 M NaCl). Amplified libraries are then obtained by adding HotStar ReadyMix 2X and primer mi× 10X and performing 12 cycles of 98°C for 15 minutes, 60°C for 10 seconds, and 72°C for 1 minute. The PCR products were further purified via AMPure XP beads. Libraries were quantified via Qubit (Thermo Fisher), and fragment sizes were analyzed via a bioanalyzer (Agilent). Libraries were paired-end sequenced via Illumina sequencing (NextSeq500, Illumina). The generated paired FastQ reads were analyzed with the cutandrun pipeline version 0.8 with default parameters and mapped to the hg19 reference genome (https://github.com/iwc-workflows/cutandrun). After filtering and duplicate removal, peaks are called via macs2. A BigWig file contains the coverage file.

### 4C-seq

4C-seq libraries were generated as previously described (16). A total of 7,5.10^6^ cells were cross-linked with 2% formaldehyde (37%) for 10 minutes. Chromatin is first digested with Csp6I (200 U, Thermo Fisher) restriction enzyme and then with DpnII (200 U, New England Biolabs). Ligation was performed with 100 U of T4 DNA Ligase (Promega) at 16°C. DNA purification was performed via NucleoMag clean-up and size selection beads (Macherey-Nagel). Inverse PCRs were performed via the Expand™ Long Template PCR System (Roche) with the primers listed in Table S2. The PCR products were quantified with a Qubit instrument (Thermo Fisher), and the fragment size was analyzed with a BioAnalyzer (Agilent). Libraries were sequenced at 75 bp single end via Illumina sequencing (MiniSeq Illumina). The generated FastQ data were analyzed with the Pipe4C version 1.1.4 pipeline (16). The data were aligned to the hg19 reference genome. A wig file containing the normalized data is available for visualization via genome browsers. Peak calling was performed with R script peakC (version 0.2) (17).

### ABC model

The activity-by-contact (ABC) prediction model is used to predict CRE promoter links (18). ABC requires an open chromatin file and an active chromatin region file. BAM files free of duplicates from ATAC-seq analysis were used to generate a fragment file, which was then converted to a tagAlign file to match the required input format. The ABC pipeline was run with the default parameters using the hg19 reference genome as the input. The data used were ATAC-seq and H3K27ac data from Capan-1 cells, with the power law used as the contact metric. The ABC score is calculated via the following equation:

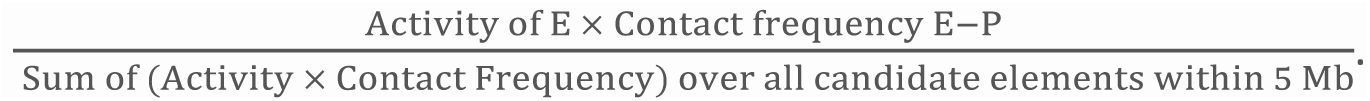

The resulting CRE-promoter prediction links were filtered to obtain only those involving the *CFTR* promoter, and a paired BED file containing the predictions was produced.

### Plasmid construction

Regions of interest were amplified via CloneAmp HiFi PCR Premix and then inserted via In-Fusion cloning (TaKaRa) into the pGL3-Basic Vector (Promega). The *CFTR* promoter (787 bp) is cloned and inserted into the HindIII restriction site, which is upstream of the firefly luciferase cDNA (*luc*). The candidate regulatory regions are subsequently cloned and inserted into the BamHI or XhoI restriction site. The PCR primers used are listed in Table S3.

CRISPR (clustered regularly interspaced short palindromic repeats) guides were designed using CRISPick and selected based on efficacy as well as predicted fidelity. Spacer sequences including compatible overhangs for cloning were ordered as single strand DNA oligos (IDT, see Table S4) and annealed in buffer B (Thermo Fisher). Annealed oligos were phosphorylated using T4 polynucleotide kinase (Thermo Fisher) and ligated into an BpiI-restricted sgRNA expression backbone (pBluescriptSKII-U6-sgRNA F+E scaffold, Addgene #74707).

### Reporter gene assay

A total of 2.5 µg of plasmid (4:1 ratio, plasmid of interest: pCMV-Bgal, internal control) was reverse transfected into 2,8.10^5^ cells with Lipofectamine 300 using 2 µL of p3000 in 12-well plates. Each condition was performed in triplicate. Seventy-two hours after transfection, the cells were lysed with 1X passive lysis buffer (Promega). The cell lysates were clarified, and 20 µL of protein extract was used for the luciferase assay and 50 µL for the beta-galactosidase colorimetric assay. Revelation reagents were purchased from Promega, and Varioskan (Thermo Fisher) was used as a plate reader. Relative luciferase activity was calculated, and Student’s t test was performed.

### CRISPR/cas9 deletion

MLV-based virus-like particles (VLP) were ordered from Leuven Viral Vector Core and produced as previously described (19). In short, 7.106 HEK293T cells were seeded in five 10 cm diameter petri dishes (BD Biosciences) and quadruple transfected with sgRNA/epegRNA/ngRNA expression plasmids, MLV-gag-pol, VSV-g and gag-cargo fusion expression constructs following the ratios described by Mangeot et al., typically withholding the BaEVRless envelope and supplementing with additional VSV-G expression plasmid using PEI (PEI, Polysciences Europe) (20). The GagMLV-Cas9 plasmid used was a kind gift of David Liu (pCMV-MMLVgag-3xNES-Cas9; Addgene #181752) (21). Supernatant containing lentiviral particles was harvested 48h and 72h post transfection, filtered through a 0.45µm pore-size filter and concentrated using centrifugation in Vivaspin columns at 3000xg. VLP productions were stored at −80°C until further use.

10 000 Capan-1 cells were incubated with the VLPs for 10 minutes and then cultured under normal cell culture conditions. After 24 hours, the medium was refreshed, and the cells were grown until sufficient numbers were obtained to perform DNA and RNA extractions. Homozygous deletion was confirmed by PCR followed by Sanger sequencing. Primers are listed in Table S4. *CFTR* gene expression was assessed by RT-qPCR using *ONEGreen* FAST qPCR Premix (Table S4).

## Results

### 1. Prediction model identifying putative regulatory elements

To identify which CRE is involved in the *cis*-regulation of the *CFTR* gene in the pancreas, we performed ATAC-seq and CUT&RUN-seq for H3K27ac in Capan-1 cells. Capan-1 cells are pancreatic duct epithelial cells derived from pancreatic adenocarcinoma that express the *CFTR* gene **(Figure 1A, Figure S1)**. ATAC-seq allows the mapping of open chromatin regions across the entire genome. H3K27ac marks are indicative of enhancer activity. We focused our interest on the *CFTR* TAD (chr7:117,039,878-117,356,812), which was separated by the CTCF boundaries at −80.1 kb and +48.9 kb, and identified multiple peaks. The active state of the *CFTR* promoter in Capan-1 cells was confirmed by the identification of ATAC and H3K27ac peaks in this region. A consensus analysis of the CUT&RUN and ATAC experiments resulted in a list of highly reliable peaks. In addition to the boundaries, six peaks were identified within the TAD. Two were located upstream of the promoter, one corresponded to the *CFTR* promoter, and three were located at the 3’ end of the *CFTR* gene **(Figure 1B)**.

**Figure 1.**
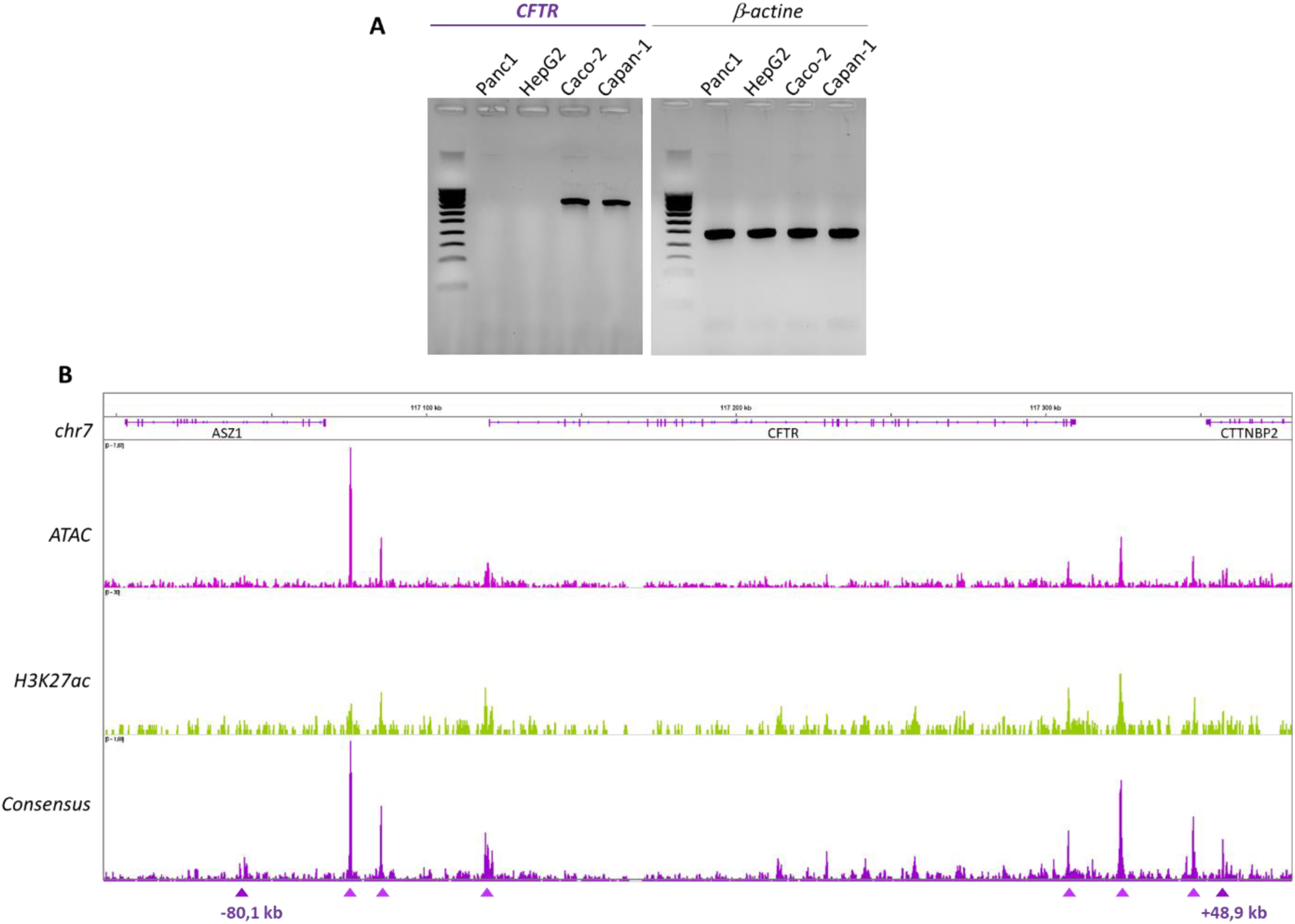
Open chromatin and enhancer mapping in Capan-1 cells. (A) RT-PCR results for CFTR expression. The CFTR gene is not express in Panc1 and HepG2 and is expressed in Caco-2 and Capan-1. On the right is the control agarose gel with B-actin expression. (B) ATAC-seq allows the mapping of open chromatin regions, i.e., regions accessible to the transcriptional machinery. We applied ATAC to Capan-1 cells and identified several peaks across the locus. A peak at the CFTR promoter was identified, indicating the accessibility of the region and confirming that CFTR is expressed in Capan-1 cells. H3K27ac is an epigenetic mark for active enhancer regions. Both marks allow cell-specific identification of the active enhancer region. A consensus analysis highlighted six major peaks as candidate regulatory regions in Capan-1 cells in addition to the TAD boundaries. The data were aligned to the hg19 genome.

ATAC and H3K27ac data enable the identification of active enhancer regions but do not provide information about the link between CREs and their target genes. To this end, we use the predictive activity-by-contact (ABC) model developed by Fulco *et al.* on the basis of a combination of experiments representing enhancer activity (ATAC-seq and H3K27ac IP) and enhancer‒promoter contact frequency (Hi-C) (18,22).

After analysis, five ABC links were highlighted, including one involving the promoter **(Figure 2)**. Three CREs were identified upstream of the promoter, two of which were known DHSs, at −44 kb and −35 kb, and one at −114 bp of the TSS (23). The downstream ABC link was mapped to the +15.6 kb DHS, previously described as an enhancer-blocking element (24,25).

**Figure 2.**
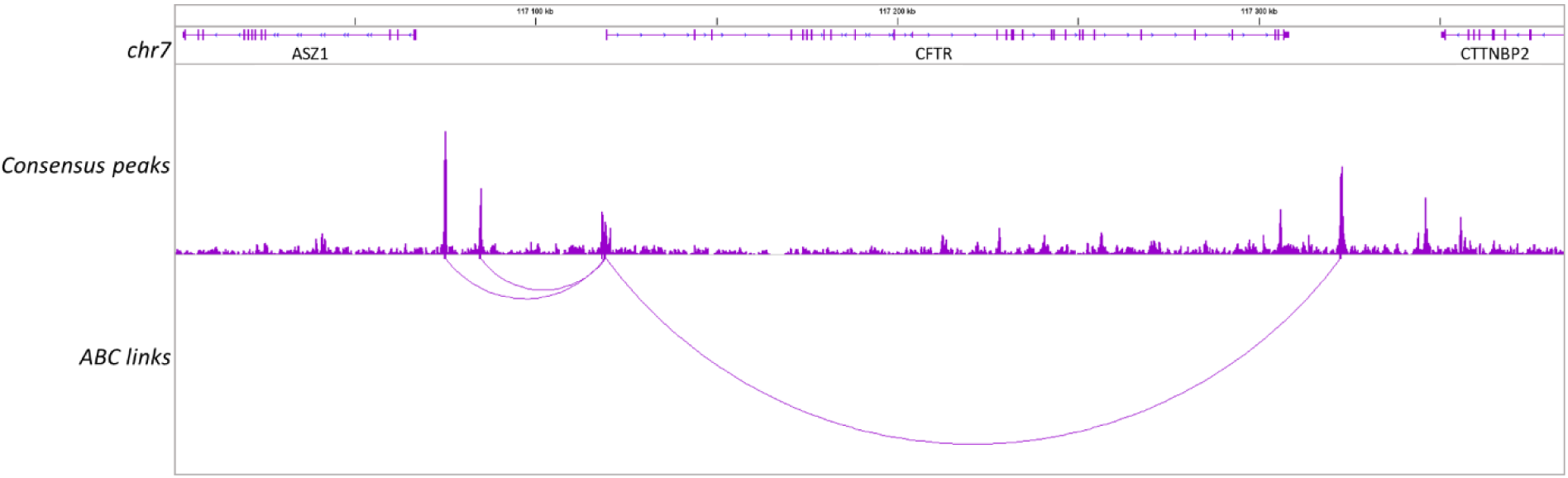
Gene–enhancer prediction links. ABC is a predictive model for identifying gene‒enhancer links via open chromatin, enhancer activity and contact frequency data. Five ABC links were identified, one of which is the promoter. Four cCREs are predicted to be involved in the regulation of the CFTR gene. The data were aligned to the hg19 genome.

### 2. Chromatin organization confirms the presence of active regulatory regions

To validate regulatory interactions, 4C-seq was applied to Capan-1 cells to determine 3D chromatin contact at the *CFTR* locus. 4C-seq, developed by the de Laat group, quantifies the frequencies of chromatin interactions with a bait of interest, for this study, the *CFTR* promoter **(Figure 3A)** (16,26). To highlight the most frequent interactions, peak calling analysis was performed on *peakC* **(Table S1)**.

**Figure 3.**
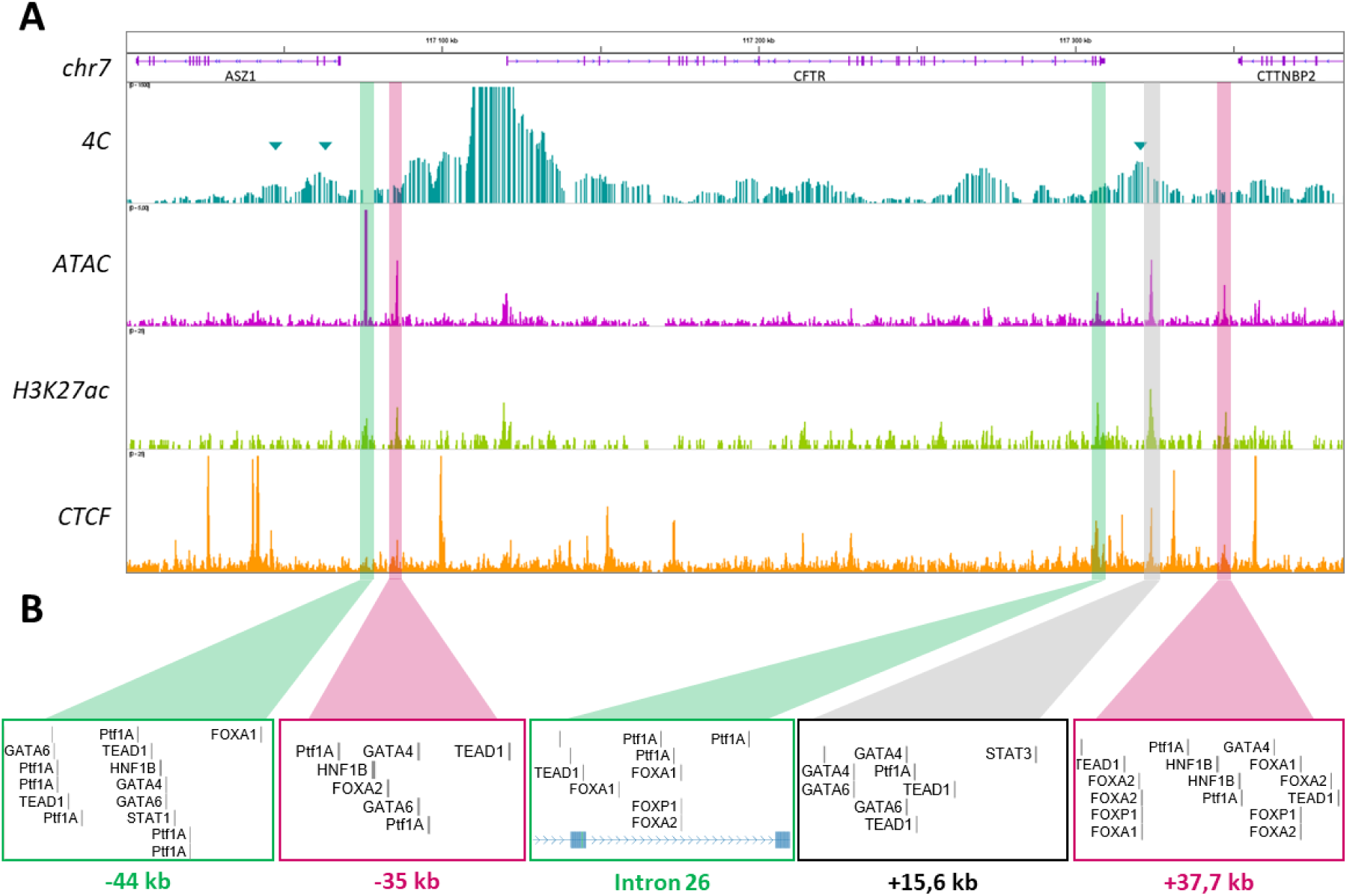
Characterization of the CFTR cis-regulatory landscape. (A) CFTR promoter interaction frequencies were determined via 4C-seq in Capan-1 cells. PeakC analyses highlighted significant interactions with the promoter (alphaFDR; 0.1). They are represented by blue arrows at the 5’ TAD boundaries −80.1 kb, at −44 kb and at +15.6 kb. ATAC-seq and CUT&RUN-seq were performed for the histone marks H3K27ac in Capan-1 cells and CTCF in Caco-2 cells. All these data were integrated to map five cCREs in detail. (B) Pancreas-specific TFs from the Jaspar 2024 TFBS were mapped to each of them (27). The data were aligned to the hg19 genome.

Among the regions identified as significant within the TAD, three were of particular interest: the −80.1 kb boundary, a region at approximately −60 kb and the +15.6 kb region **(Figure 3A, blue arrows)**. The interaction with the +15.6 kb region was confirmed to be highly important, in alignment with the predictions of the ABC model. However, the −44 kb and −35 kb regions identified in the ABC model do not overlap with a significant 4C peak. Nevertheless, there is a significant interaction between the promoter and upstream region, which can bring the −44 kb and −35 kb regions in proximity to the promoter. A DHS within the last intron, intron 26, of the *CFTR* gene corresponded to an H3K27ac peak as well as a 4C peak. The same observations were made for a region at +37.7 kb, which is near a previous DHS identified in the lung, the +36.6 kb region (23). On the basis of this information, five cCREs were delineated. Following transcription factor (TF) alignment, putative exocrine pancreas-specific TF binding sites were confirmed in each of the five cCREs **(Figure 3B)**.

### 3. Luciferase assays characterize cCRE activity

To evaluate the activity and function of the candidate regulatory regions involved in *CFTR* regulation, *in vitro* luciferase assays were conducted. The initial analysis focused on the cCREs identified by the chromatin assay and the ABC model. The regions of interest were cloned and inserted into the pGL3-Basic vector, in which the minimal *CFTR* promoter (787 bp, chr7:117,119,328-117,120,114, hg19) drives the expression of the luciferase gene (LUC). Each construct was co-transfected into Capan-1 cells with a pCMV-beta-galactosidase control plasmid. Luciferase activity significantly increased in the presence of the −44 kb region (1.7-fold) and the DHS in intron 26 (1.8-fold) **(Figure 4)**. Compared with the promoter alone, +15.6 kb did not affect the luciferase activity. The −35 kb and +37.7 kb regions had a minor silencing effect (−1.39-fold).

**Figure 4.**
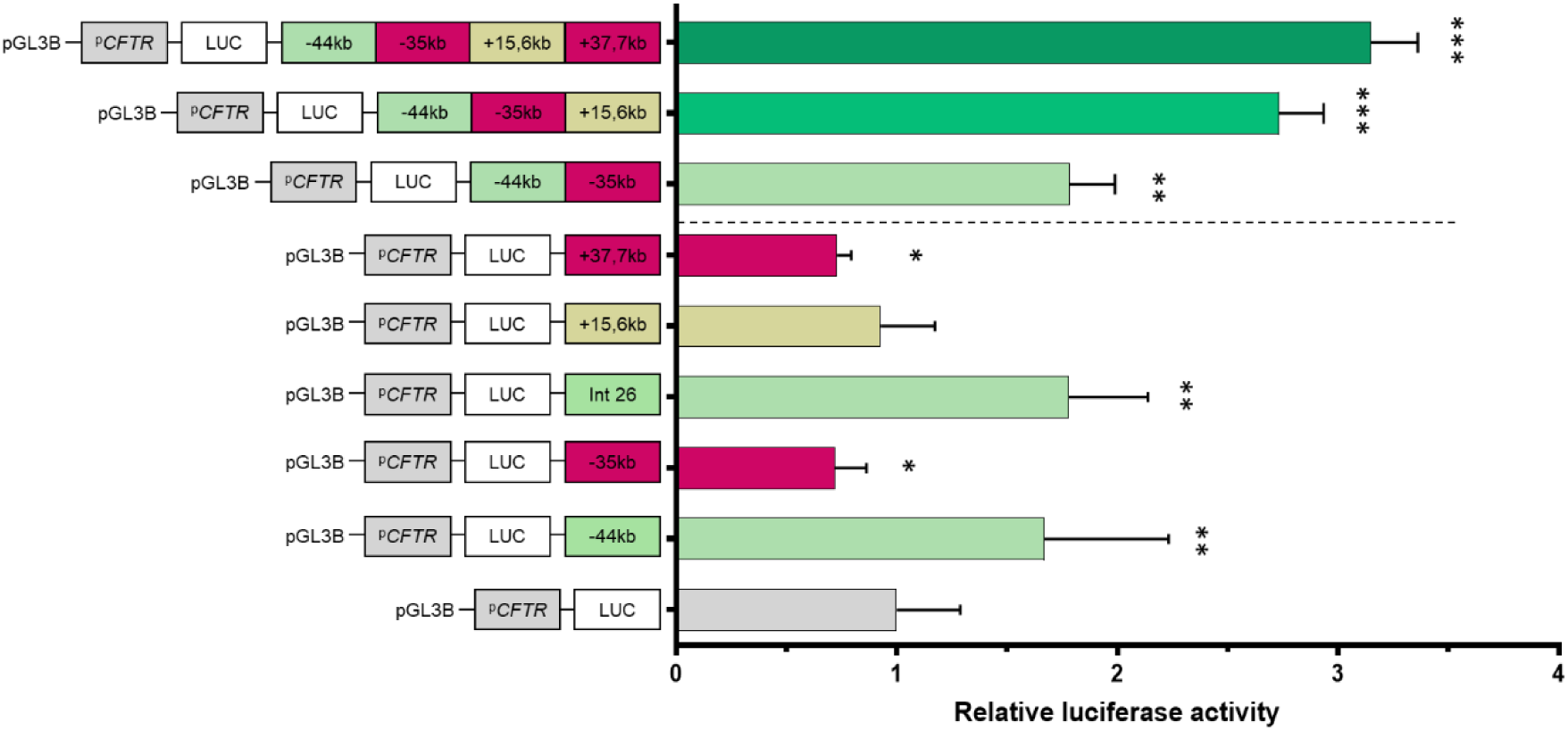
Activity assay of identified pancreatic-specific cCREs. Capan-1 cells were transfected with luciferase reporter constructs containing the CFTR basal promoter (PCFTR; 787 bp) and cCREs identified via chromatin analysis. Two regions, −44 kb and DHS in intron 26, show enhancer activity, whereas −35 kb and +37.7 kb show slight decreases in activity. +15.6 kb shows no effect. The luciferase data are shown relative to the CFTR basal promoter vector (set to 1). The error bars represent the standard deviation (SD; n = 9), *<0,05;**<0,001;***<1.10^—9^ using unpaired t-tests.

Although the single CRE luciferase reporter data is informative, it is important to consider the fact that in the genome, there are simultaneous promoter–CRE links; this is not an individual relationship. Several studies have shown that gene expression can be affected by one or multiple CREs (28–30). Indeed, *cis*-regulation is a process whereby multiple CREs are involved in the formation of chromatin modules that in turn regulate the same gene (31). To evaluate the cooperative effect of the identified cCREs that share putative TF binding sites, we designed three synthetic constructs: one encompassing the −44 kb and −35 kb regions; another containing the −44 kb, −35 kb and +15.6 kb regions; and the last one with the four cCREs, −44 kb, −35 kb, +15.6 kb and +37.7 kb. The combination of two cCREs was observed to induce an increase in luciferase activity, although this increase was not significantly different from the effect observed with the −44 kb region alone **(Figure 4).** When the +15.6 kb region is added, a 2.7-fold increase is measured, and with the combination of the four regions, we observe a 3.2-fold increase. There is a synergistic effect of the CREs.

To extend the functional analysis of the regulatory elements in pancreatic cells, we also tested regions that have been described as cCREs in previous studies (32,33). Four regions were analyzed: −3.4 kb and DHSs in introns 11, 18 and 23. The −3.4 kb region has a moderate enhancing effect on pulmonary cells (34); intron 11 is described as an enhancer in intestinal cells, and chromatin interactions have been shown in pancreatic cell lines (33,35). DHSs have been identified in introns 18 and 23, and specific TF binding has been shown in intron 18 (32). In our hand, the DHS in intron 18 had no effect on the luciferase assay (**Figure 5A**). The −3.4 kb region and intron 11 seem to have a small enhancer effect, whereas intron 23 shows highly significant enhancer activity (3.5-fold). Alignment with specific exocrine pancreas TFs revealed the presence of an HNF1B motif in all cCREs, except for intron 18, which showed no effect in the luciferase assay (**Figure 5B**).

**Figure 5.**
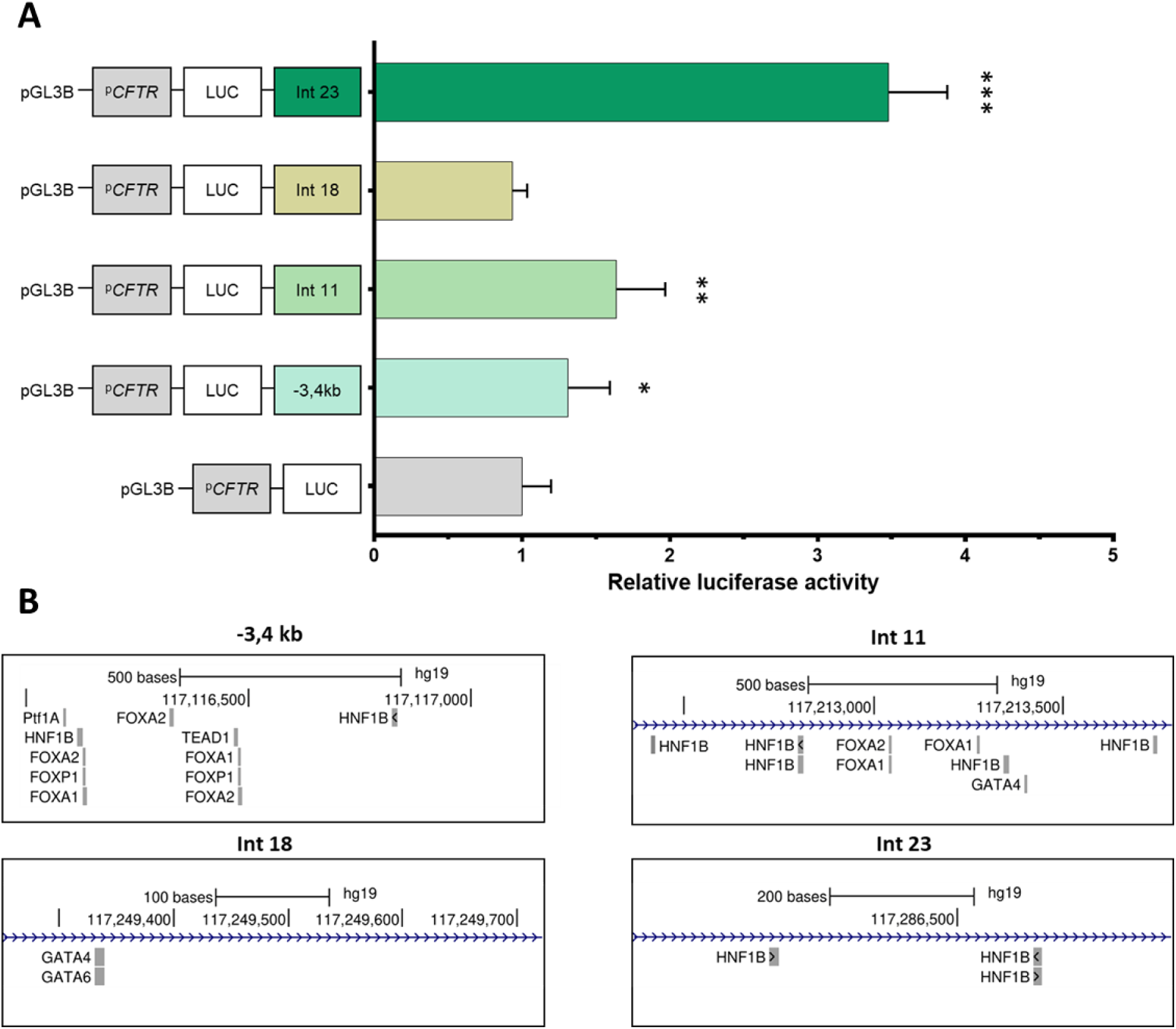
Activity assay on previously described cCREs. (A) Capan-1 cells were transfected with luciferase reporter constructs containing the CFTR basal promoter (PCFTR; 787 bp) and cCREs described in the literature. We confirmed the enhancer effect of regions −3,4 kb, the DHS in intron 11 and intron 23, with a major effect on the DHS in intron 23. The luciferase data are shown relative to the CFTR basal promoter vector (set to 1). The error bars represent the standard deviation (SD; n = 9), *0,05;**<0,0001;***<1.10^−8^ using unpaired t-tests. (B) Pancreatic-specific TFs from the Jaspar 2024 TFBS were mapped.

Finally, five enhancers have been described: the region at −44 kb, at −3.4 kb and the DHS within introns 11, 23 and 26. Two regions have minor silencing effects when present alone: the region at −35 kb and at +37.7 kb. Cooperative effects have been observed with the combination of CREs, with an important increase in activity.

### 4. Endogenous assay validated the enhancer effect of the −44 kb region

To confirm our observations based on chromatin structural assays (ATAC-seq, CUT&RUN-seq and 4C) and experimental quantification using the reporter assay, we performed precise deletion of the endogenous −44 kb enhancer region using CRISPR/Cas9 in Capan1 cells. Two flanking single guide RNAs (sgRNAs) were designed and positioned at each extremity of the targeted region. To achieve homozygous deletion, efficient delivery of the genome editing technology is essential. For that reason, virus-like particles (VLPs) were produced containing the −44kb flanking sgRNAs **(Figure 6A)**. Homozygous deletion was confirmed by PCR followed by Sanger sequencing, validating a clone with a 2166 bp deletion **(Figure 6A, B)**. The impact of the CRE deletion on *CFTR* gene expression was assessed by RT-qPCR and compared to untreated Capan-1 control cells. The deletion of the −44 kb region resulted in a reduction of *CFTR* expression to 15% of that in untreated cells **(Figure 6C)**. These findings are consistent with the results of the luciferase reporter assay, which demonstrated a slight increase in activity in the presence of the −44 kb CRE. **(Figure 4)**. These results demonstrate that the combination of ATAC-seq, CUT&RUN-seq, 4C and the luciferase reporter assay allows to identify and validate new enhancer regions.

**Figure 6.**
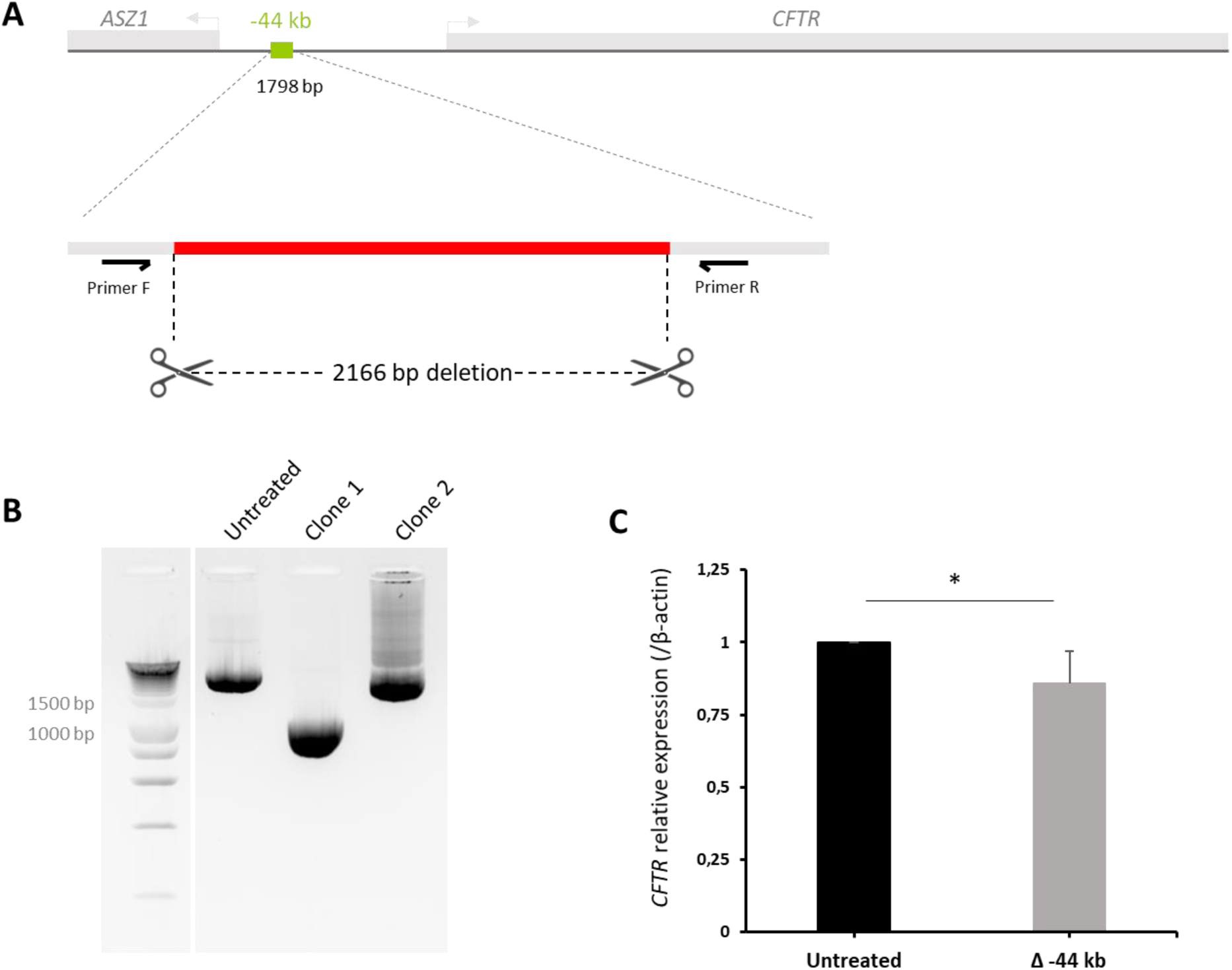
Endogenous effect of CRE deletion. (A) The 1798 bp of the −44 kb enhancer region was deleted by positioning two sgRNAs at each extremity. After RT-PCR and Sanger sequencing we validated a homozygous deletion of 2166 bp showed in (B). Results of RT-PCR with primers positioned outside the deleted region. Amplifications were performed on untreated cells and on the selected clone. PCR products from untreated cells are approximately 3000 bp (3239 bp). PCR products from Clone 1 are approximately 900 bp, consistent with Sanger sequencing results showing a deletion of 2166 bp. Clone 2 is negative, showing the same size product as the control condition. (C) RT-qPCR was performed on Clone 1, revealing a slight decrease (15%) in CFTR gene expression normalized to the B-actin gene, compared to the untreated condition. Results are based on two technical replicates and three biological replicates. *<0,05 using unpaired t-tests.

### 5. Exploring the landscape outside the TAD

Although many CRE-promoter interactions occur within the same TAD, it has been shown that some CREs can interact with promoters in different TADs, which is called boundary stacking (36). Taking this into account, we extended our analysis of chromatin data beyond the scope of the *CFTR* TAD (**Figure 7A**). Peak calling analysis of the 4C-seq data revealed a region of interest near the *LSM8* gene. The promoter of the *CFTR* gene significantly interacts with a region located at +485 kb from the last codon of the *CFTR* gene. To gain insight into the functional role of this region, which was mapped to 4C, ATAC-seq, H3K27ac and multiple CTCF peaks, the region was aligned to the ENCODE database (https://screen.encodeproject.org/) **(Figure 7B)**.

**Figure 7.**
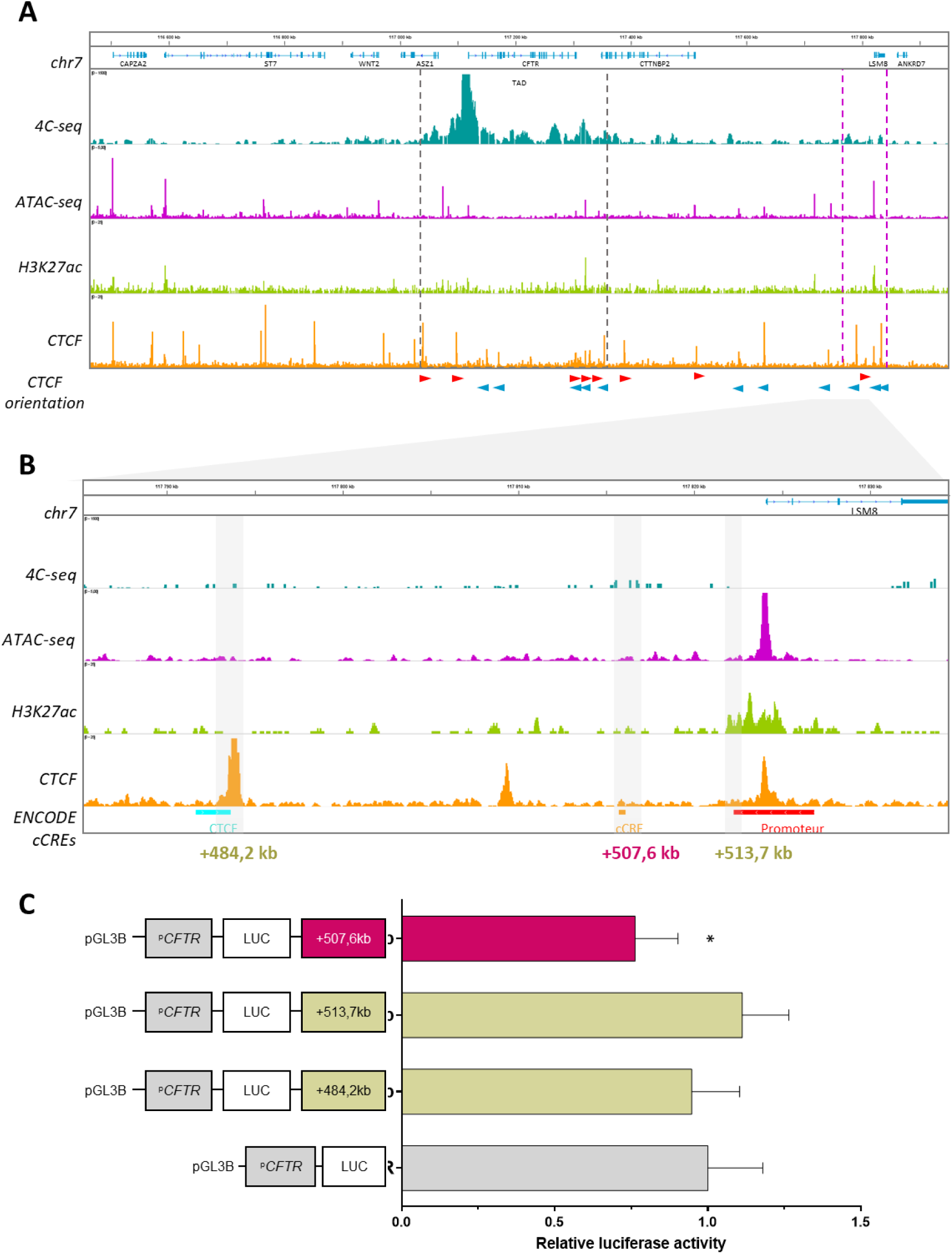
Exploration of regulatory elements beyond TAD boundaries. (A) Frequencies of CFTR promoter interactions were determined by 4C-seq in Capan-1 cells. ATAC-seq and CUT&RUN-seq were performed for the histone marks H3K27ac in Capan-1 cells and CTCF in Caco-2 cells. The orientation of each CTCF site is indicated by a blue or a red arrow. A region of interest has been selected in pink and zoomed in (B). The ENCODE cCREs database indicated three putative functional elements, a CTCF site in blue, an enhancer-like site in orange and a promoter-like site in red. (C) These three regions were selected for use in luciferase assays. Capan-1 cells were transfected with luciferase reporter constructs containing the CFTR basal promoter (PCFTR; 787 bp) and cCREs. We observed a silencer effect in the +507.6 kb region. The luciferase data are shown relative to the CFTR basal promoter vector (set to 1). The error bars represent the standard deviation (SD; n = 9), *<0,05 using unpaired t-tests. The data were aligned to the hg19 genome.

Analysis of the 4C data revealed that the higher signal region mapped to a CTCF peak at +484.2 kb and to a peak at +507.6 kb. Mapping to the ENCODE cCREs database showed that it corresponds to a predicted CRE region. This region also encompasses the promoter of the *LSM8* gene, which has an active chromatin signal in our data. Using these elements, we selected these three regions for functional assay testing **(Figure 7C)**. They were subsequently cloned and inserted into the pGL3-Basic vector with the minimal *CFTR* promoter, and a luciferase assay was then performed. As expected, the CTCF site had no effect on the observed activity. A similar observation was made for the +513.7 kb region corresponding to the *LSM8* promoter. However, for the +507.6 kb region, which corresponds to the predicted CRE region, a silencer effect was observed (−1.32-fold). Those preliminary results demonstrate that while most CRE-promoter interactions occur within the same TAD, interactions can also extend beyond TAD boundaries and emphasizing the importance of exploring inter-TAD interactions for a comprehensive understanding of gene regulation.

### 6. Cell type specificity

To gain further insights into tissue-specific regulation of *CFTR* expression, we intended to compare data from other cell types. This allows to identify tissue-specific characteristics. ATAC-seq data was generated for the intestinal Caco-2 cell line, and publicly available data for HepG2 cells were retrieved (14) **(Figure 8A)**. Caco-2 cells are derived from a colorectal adenocarcinoma and express the *CFTR* gene, whereas HepG2 cells are derived from hepatocellular carcinoma and do not express the *CFTR* gene **(Figure 1A)**.

**Figure 8.**
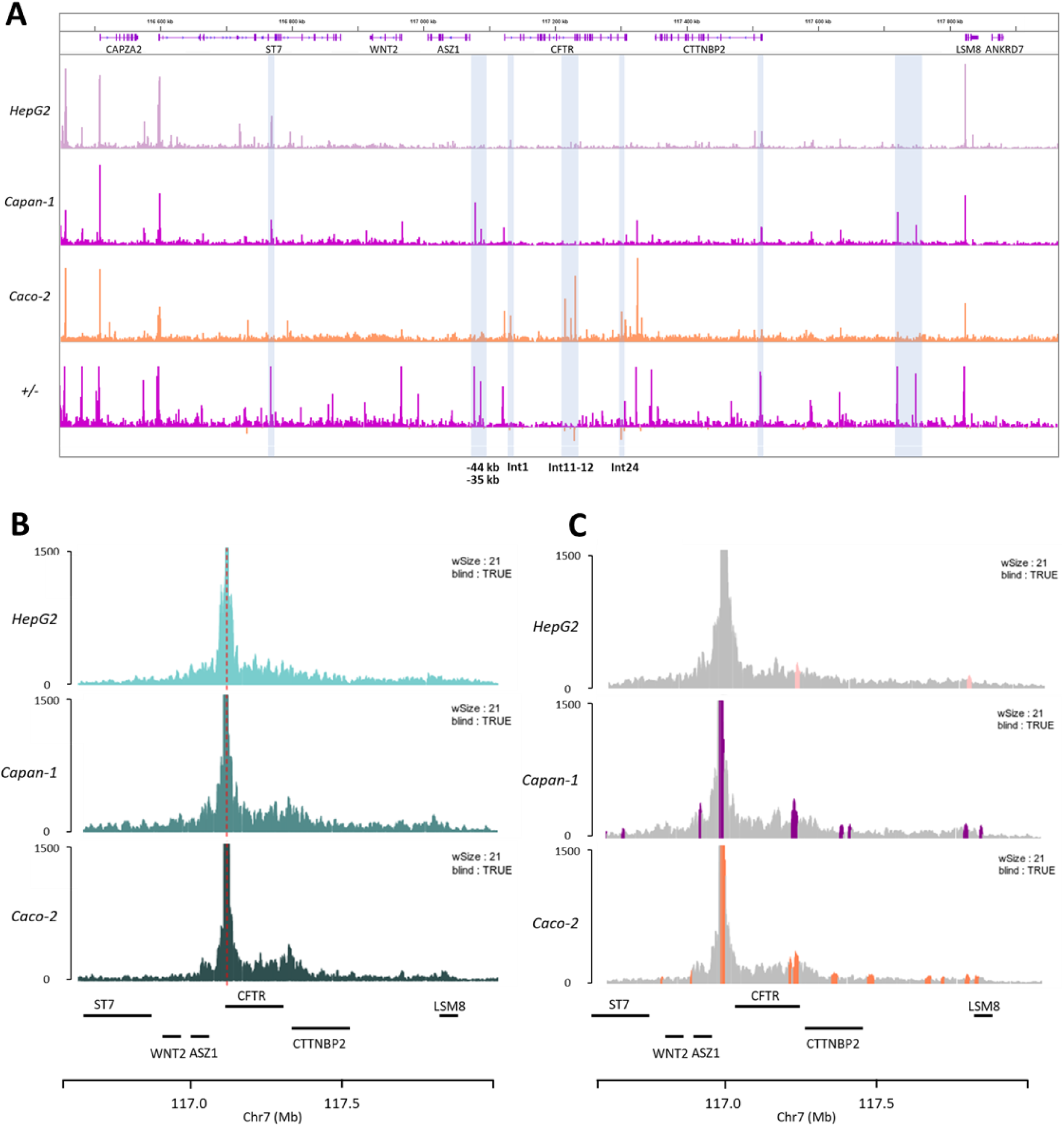
Tissue specificity of cis-regulation. (A) ATAC-seq data for Capan-1, Caco-2 and HepG2 cells to compare the tissue specificity of DHSs. HepG2 cells do not express the CFTR gene, and any DHS is detected at the CFTR locus. BigWig comparison (+/−) of Capan-1 and Caco-2 data revealed few differences (highlighted in blue), especially at the CFTR locus. −44 kb and −35 kb are specific for Capan-1 cells, and introns 1, 11, 12 and 24 **are specific** for Caco-2 cells. (B) CFTR promoter interaction frequencies were determined by 4C-seq in Capan-1, Caco-2 and HepG2 cells. HepG2 cells show only random interactions, whereas Capan-1 and Caco-2 cells show a specific profile. (C) Significant regions are highlighted by colored peaks with shared and unique regions (alphaFDR; 0,4). The data were aligned to the hg19 genome.

As expected, a comparison of open chromatin data profiles from *CFTR*-expressing cells and a cell line that does not express the gene revealed a notable difference in the profiles **(Figure 8A)**. In HepG2 cells, no significant signal was observed around the *CFTR* locus. In the case of the *CFTR*-expressing cells, we observed a greater degree of similarity within the TAD, although interestingly, there were also some differences. A BigWig comparison analysis **(Figure 8A, +/−)** between Capan-1 and Caco-2 cells revealed some significant differences. The DHS signal in Caco-2 cells is more pronounced within intronic regions of the *CFTR* gene, confirming previous data that identified enhancers within introns 1, 11, 12, 24 and 26 (35). For the Capan-1 cells, the majority of the peaks were observed outside the *CFTR* gene. Notably, there was a significant disparity between the two cell lines, with the −44 kb and *CFTR* 3’ regions lacking DHS peaks in Caco-2 cells. This finding appears to be specific to the pancreas. Differences were also observed in the DHS peaks outside the TAD. For example, the peaks located between the *CTTNBP2* and *LSM8* genes are exclusive to Capan-1 cells.

4C-seq data from Caco-2 and HepG2 cells **(Figure 8B)** evidences a clear difference between cells expressing the *CFTR* gene and HepG2 cells, where there are more random interactions and they decrease as we move away from the promoter. The profiles of Capan1 and Caco-2 cells appear to be similar. PeakC analysis was conducted on both datasets, revealing identical interactions as well as some distinct interactions. For example, a significant peak encompassing intron 24, which is absent in Capan-1 cells, is observed in Caco-2 cells **(Figure 8C).** We show with ATAC-seq data that the major difference in the open chromatin regions between pancreatic and intestinal cells is in the cluster of introns 11 and 12 and within intron 24. A previous publication demonstrated that the DHS in intron 12 has a significant enhancing effect and that the combination of the DHSs in introns 12 and 24 has a notable effect on Caco-2 cells (35). We repeated the luciferase reporter assay in both cell types **(Figure 9A)**. In Caco-2 cells, intron 12 increased luciferase activity by 15-fold, and the combination of intron 12 and intron 24 increased luciferase activity by 50-fold. In contrast, no increase in luciferase activity was observed in Capan-1 cells in response to either of these regions. These enhancers hence appear specific to intestinal cells.

**Figure 9.**
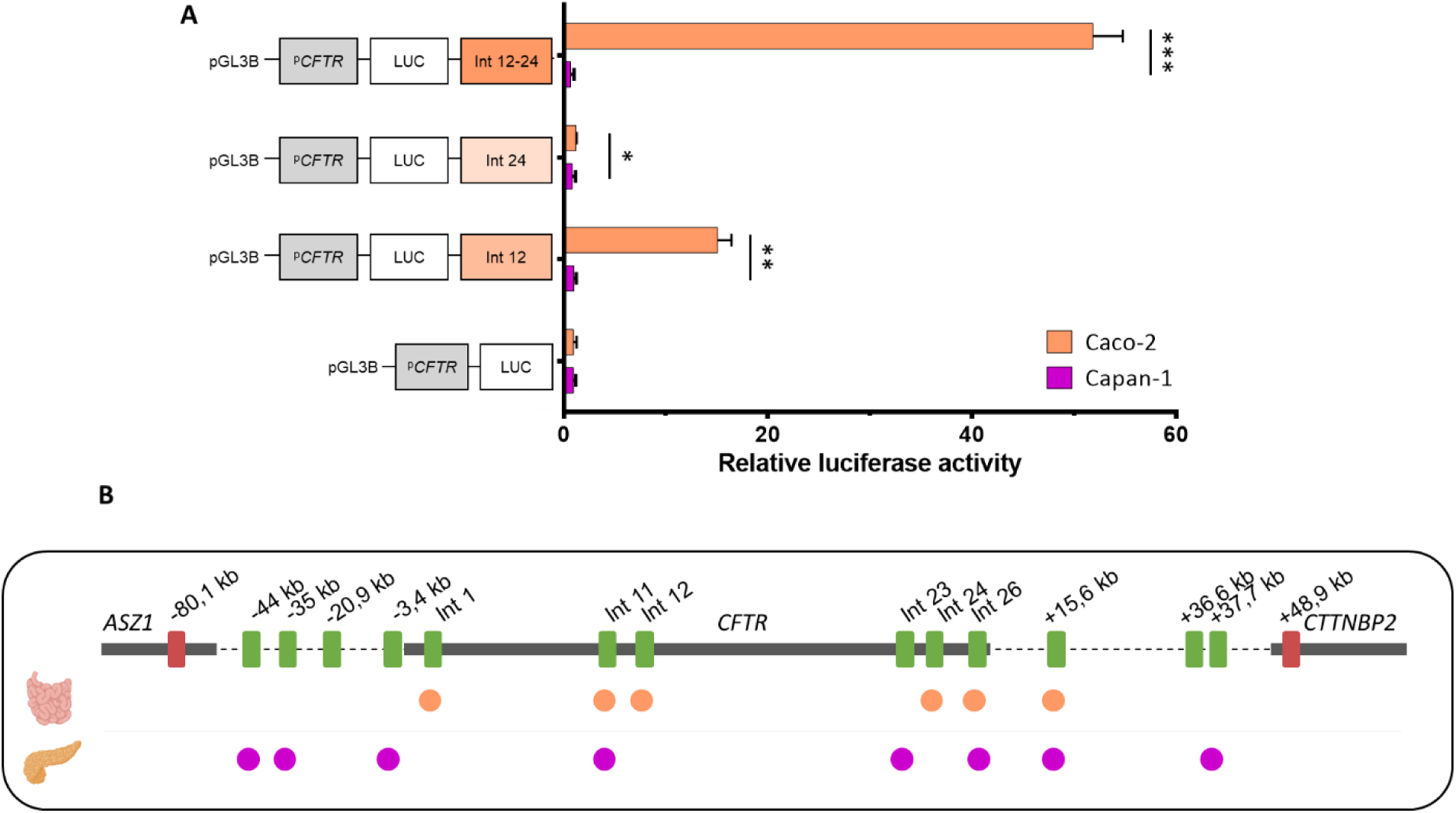
Activity of regulatory regions in Caco-2 and Capan-1 cells. (A) Capan-1 and Caco-2 cells were transfected with luciferase reporter constructs containing the CFTR basal promoter (PCFTR; 787 bp) and CREs. We observed an enhancing effect in Caco-2 cells for introns 12 and 24 and the combination, whereas no effect was detected in Capan-1 cells. The luciferase data are shown relative to the CFTR basal promoter vector (set to 1). The error bars represent the standard deviation (SD; n = 9), *<0,001;**<1.10^−15^;***<1.10^−19^ using unpaired t-tests. (B) Linear view of the CREs involved in CFTR regulation in the small intestine, with a higher proportion of intronic CREs. In the pancreatic cells, active CREs are rather present upstream the CFTR gene as well as in the last introns of the gene and downstream.

These chromatin analyses revealed that the *cis*-regulatory elements involved in the long-distance regulation of the *CFTR* gene exhibit a degree of tissue specificity, as illustrated in **Figure 9B**.

## Discussion

To gain insight into the regulatory landscape of the *CFTR* gene, we set out to generate a 3D genome characterization in pancreatic cells. The *CFTR* gene has complex and incompletely described gene regulation at the temporal and tissue scales, and distal regulatory elements have been shown to regulate gene expression (12). The best-described long-range regulatory mechanism involves the lung and intestine, but there is a considerable lack of long-range regulatory mechanisms in the pancreas, one of the first organs to display temporal CF-related symptoms early in life (even *in utero*).

Importantly, there is a lack of relevant pancreatic models that express the *CFTR* gene; for example, the widely used Panc1 cell line does not express the *CFTR* gene. Consequently, there is a lack of available chromatin data. To resolve this, we generated ATAC, H3K27ac and 4C data, in Capan-1 cells, a cell line of which we demonstrate *CFTR* gene expression.

For informative genome conformation studies, it is crucial to integrate multiple, complementary assays. The ATAC assay provides information on accessible DNA regions; however, these data must be contextualized with histone mark mappings to determine the functional role of the region. 4C provides insight into chromatin interactions. However, it is important to consider that not all interactions necessarily lead to transcriptional activation (37). By combining these approaches, we identified potential regulatory regions. It remains however important to investigate whether all the active regions surrounding the *CFTR* gene are truly involved in its regulation. This step, which is crucial for determining the target gene, represents a significant challenge in understanding long-range regulation.

To address this issue, we decided to use the ABC predictive model (18). For several years, numerous computational methods for determining gene‒enhancer links have been developed and are still being developed. The ABC model is widely used because of the ease of use made by developers, and it has also been validated by users (38–41). Nonetheless, the ABC model uses ATAC, H3K27ac and Hi-C data and thus does not identify other CREs as silencers or Epromoters, which are promoters with an enhancer function (42). The predictive model helps us to prioritize the candidate regions. Four distal enhancers are predicted to regulate the *CFTR* gene; indeed, in this model, each gene is regulated by an average of 2.8 enhancers (22).

The 4C data provide complementary information on the 3D interactions of the *CFTR* promoter to validate our candidate regulatory regions, as recommended in a previous publication (3). The +15.6 kb region is a known DHS previously described in pulmonary and intestinal cells as an enhancer-blocking region (25). Figure S2 shows that in Caco-2 cells, enhancer blocking limits the propagation of H3K27ac. In Capan-1 cells, we confirmed the presence of a DHS, which is marked by H3K27ac and interacts with the *CFTR* promoter. By definition, an enhancer blocking is located between an enhancer and the protected promoter. In the case of the +15.6 kb region, we observed a small ATAC and H3K27ac just downstream, representing the +37.7 kb region, which could be an enhancer blocked by the +15.6 kb region. In the native context, in the presence of the +15.6 kb region, the +37.7 kb region does not have an enhancer effect and slightly reduces the activity in the luciferase assay.

The −44 kb and −35 kb regions predicted to target the *CFTR* gene correspond to both H3K27ac and ATAC peaks, but no major interaction was observed in 4C data. The −44 kb region shows an enhancer effect but not the −35 kb region, which has a similar effect to that of the +37.7 kb region. From previous studies, we know that there is also an enhancer-blocking upstream of the promoter at −20.9 kb, which corresponds to a 4C peak in Capan-1 cells but not an ATAC peak (25,43). It remains to be determined whether this cCRE is active in pancreatic cells, as the CTCF data are from Caco-2 cells. The last cCRE identified by chromatin data is located in intron 26 and has not been identified by the ABC model, but it corresponds to the 4C, ATAC, and H3K27ac peaks and has increased activity by 1.8-fold in the luciferase assay.

Because *cis*-regulation is hosted by a chromatin module, we wanted to add information about TFs to obtain a complete picture of this landscape (31). We aligned TFs from JASPAR, as there is no ChIP-seq data for whole-genome exocrine pancreas-specific TFs in Capan-1 cells. It would be of interest to perform ChIP with HNF1B, Ptf1a and FOXA1/2, as these proteins are more common in the active CRE. The STRING database (https://string-db.org/) revealed that HNF1B and Ptf1a are associated with one another **(Figure S3)**. On the basis of the chromatin module, we also provide information on the cooperative effect of the CRE (28). Indeed, the combination of CREs has a synergistic effect. The activity of each individual enhancer is lower than that of the combination (30). Using Cas9-based genome engineering efficiently delivered through VLPs, we were able to confirm the role of the −44kb region in an endogenous setting. Although we demonstrate an endogenous impact of the individual deletion of the −44 kb region, considering the importance of the chromatin module, it would be interesting to go further by individually deleting the other CREs, as well as performing multiple deletions to determine whether the impact on *CFTR* expression is pronounced than what we observe here. Subsequently, chromatin states will need to be assessed to understand the consequences of CRE loss.

The majority of CRE-gene interactions are located within 100 - 500 kb of the TSS (18,44). Hence, we wanted to extend our characterization beyond the TAD. Interestingly, we identified a significant long-range interaction 500 kb downstream of the TSS. By observing the orientation of the CTCF sites and based on the processed loop extrusion, we noticed that this region lies within the neighboring chromatin loop (45). Studies have demonstrated the possibility of cross-TAD interactions, notably through boundary stacking (36). Therefore, characterizing cCREs outside the TAD would be interesting. In the luciferase reporter assay, we also measured the silencer effect of this +507.6 kb region, which was predicted to be functional by the ENCODE database. In the database, it was annotated as an enhancer, but it is important to note that there is no silencer category. They are less well-characterized than enhancers, and we must explore the binding of repressive TFs while considering that the luciferase assay performed is far from the native context. Additionally, it will be necessary to explore the region using technologies, such as those developed by Brosh *et al*., that account for larger DNA sequences, including both the target gene and its regulatory elements (46).

Importantly, *cis*-regulation is a tissue-specific process, and the 3D chromatin architecture changes depending on the cell type. In HepG2 cells, DHS and 4C signals are absent, whereas in Caco-2 and Capan-1 cells, chromatin architecture is important for regulating *CFTR* gene expression. Notably, only 19% of enhancer–gene pairs are shared across distinct cell types (22). Our observations are consistent with this hypothesis, as the CREs identified in Caco-2 cells are not present in Capan-1 cells. It is important to note that our data were obtained from Capan-1 cell lines. Further studies are needed using more relevant models to strengthen our findings. Primary cells would be ideal, although they are much more challenging to work with and engineer (47).

## Conclusions

In summary, we have provided chromatin genome-wide data in pancreatic Capan-1 cells, and our focus on the *CFTR* locus provides novel insights on the distal regulation of the *CFTR* gene with the identification of several CREs as illustrated in **Figure 10**. In light of the increasing interest in non-coding variations in complex genetic diseases, a deeper understanding of chromatin architecture is crucial for advancing our knowledge in this field.

**Figure 10.**
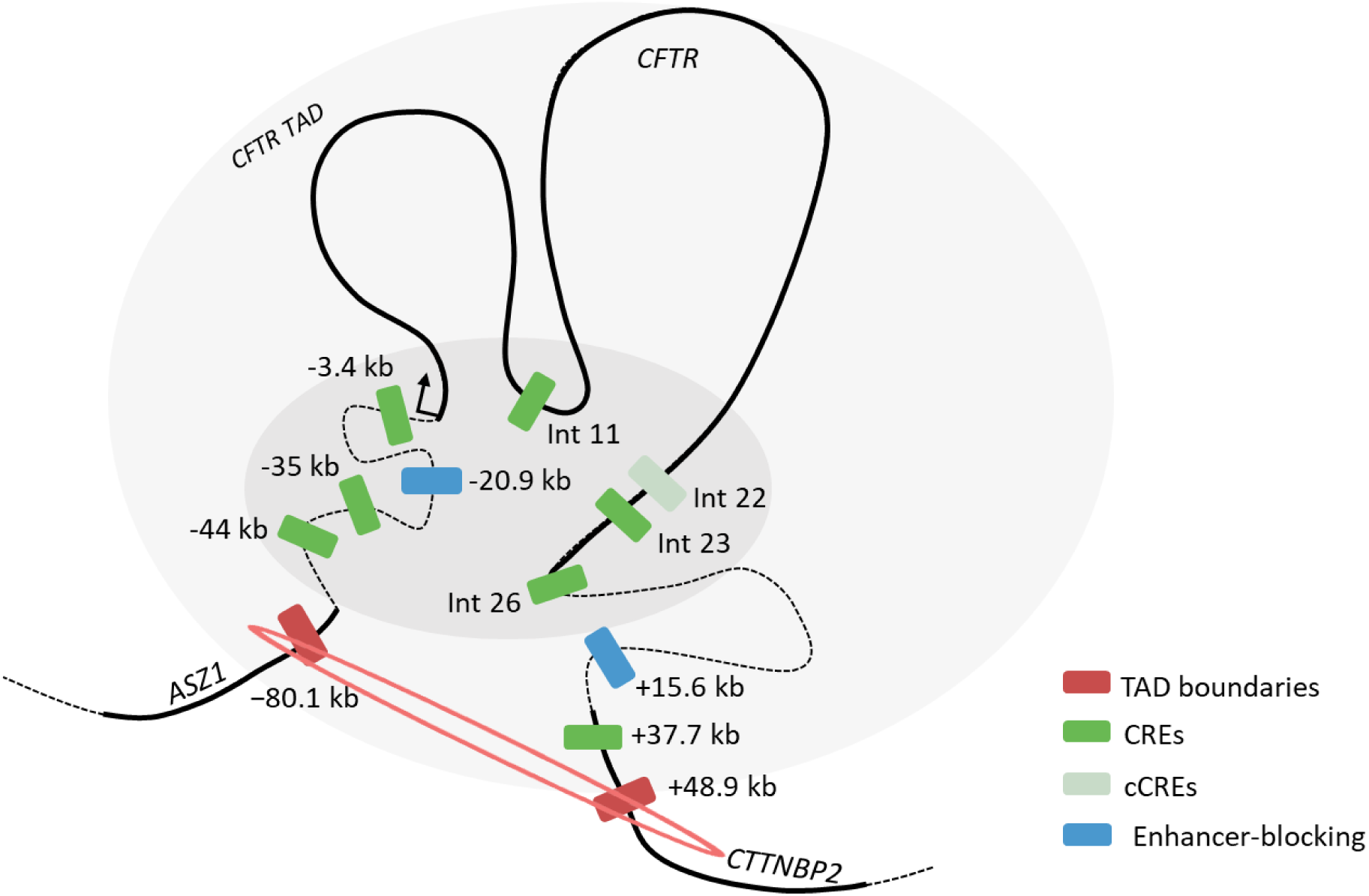
Schematic overview of the CFTR regulatory landscape. We provide a predictive three-dimensional model of CFTR regulation in pancreatic cells. Chromatin loops are represented with black lines with coding sequences as thick lines. TAD boundaries are highlighted in red, candidate CRE in light green and CREs in green. Enhancer-blocking elements are shown in blue.

## List of abbreviations

4C: circular chromosome conformation capture
ABC: activity-by-contact
ATAC: assay for transposase-accessible chromatin
CBAVD: congenital bilateral absence of vas deferens
CF: cystic fibrosis
CFTR: *cystic fibrosis transmembrane conductance regulator*
CFTR-RD: CFTR-related diseases
cCRE: candidate *cis*-regulatory elements
CRE: *cis*-regulatory elements
CRISPR: clustered regularly interspaced short palindromic repeats
CTCF: CCCTC-binding factor
DHS: DNase I hypersensitive site
CUT&RUN: Cleavage Under Targets & Release Using Nuclease
gRNA: guide RNA
PTM: post-translational modifications
TAD: topologically associating domain
TF: transcription factor
TSS: transcription start site

## Declarations

### Ethics approval and consent to participate

Not applicable.

### Availability of data and materials

Sequencing data from this study are available at NCBI GEO under accession number GSE284199, GSE284200, GSE284414.

### Competing interests

The authors declare that they have no competing interests.

### Funding

This work was supported by grants from the French foundation “Vaincre la mucoviscidose”, the “ministère de l’Enseignement supérieur et de la recherche”, and the associations “Gaétan Saleün” and “PLB Muco”.

Cas9-VLP work was funded by grants from the Mucovereniging Belgium and Fund Alphonse Jean Forton from the King Baudouin Foundation (2020-J1810150-E015), Emily’s Entourage (US_172749169_6). M.B. was supported by an FWO-SB doctoral fellowship 1SE8122N.

### Author contributions

Conceptualization, C.B., S.M.; methodology, investigation, software, validation, and formal analysis, C.B.; investigation for ATAC and CUT&RUN, A.L.N., investigation for Caco-2 luciferase assay, M.C., Cas9-VLP design and production, M.B., supervision for CRISPR/cas9, M.S.C.; writing—original draft preparation, C.B.; writing—review and editing, C.B., S.M., A.L.N, M.B., M.S.C.; funding acquisition, S.M., C.F, M.S.C. All authors have read and agreed to the published version of the manuscript.

## Acknowledgements

We acknowledge the Leuven Viral Vector Core for the production of VLPs.

## Notes

### Competing Interest Statement

The authors have declared no competing interest.

## References

1. The ENCODE Project Consortium, Abascal F, Acosta R, Addleman NJ, Adrian J, Afzal V, et al. Perspectives on ENCODE. Nature. 2020 Jul 30;583(7818):693–8.

2. Moore JE, Purcaro MJ, Pratt HE, Epstein CB, Shoresh N, Adrian J, et al. Expanded encyclopaedias of DNA elements in the human and mouse genomes. Nature. 2020 Jul;583(7818):699–710.

3. Gasperini M, Tome JM, Shendure J. Towards a comprehensive catalogue of validated and target-linked human enhancers. Nat Rev Genet. 2020 May;21(5):292–310.

4. Lopez Soriano V, Dueñas Rey A, Mukherjee R, Inglehearn CF, Coppieters F, Bauwens M, et al. Multi-omics analysis in human retina uncovers ultraconserved cis-regulatory elements at rare eye disease loci. Nat Commun. 2024 Feb 21;15(1):1600.

5. Zaugg JB, Sahlén P, Andersson R, Alberich-Jorda M, de Laat W, Deplancke B, et al. Current challenges in understanding the role of enhancers in disease. Nat Struct Mol Biol. 2022 Dec;29(12):1148–58.

6. The 100,000 Genomes Project Pilot Investigators. 100,000 Genomes Pilot on Rare-Disease Diagnosis in Health Care — Preliminary Report. New England Journal of Medicine. 2021 Nov 10;385(20):1868–80.

7. Pang B, van Weerd JH, Hamoen FL, Snyder MP. Identification of non-coding silencer elements and their regulation of gene expression. Nat Rev Mol Cell Biol. 2023 Jun;24(6):383–95.

8. El-Seedy A, Ladeveze V. CFTR complex alleles and phenotypic variability in cystic fibrosis disease. Cell Mol Biol. 2024 Sep 8;70(8):244–60.

9. Sermet-Gaudelus I, Girodon E, Vermeulen F, Solomon GM, Melotti P, Graeber SY, et al. ECFS standards of care on CFTR-related disorders: Diagnostic criteria of CFTR dysfunction. Journal of Cystic Fibrosis. 2022 Nov 1;21(6):922–36.

10. Simmonds NJ, Southern KW, De Wachter E, De Boeck K, Bodewes F, Mainz JG, et al. ECFS standards of care on CFTR-related disorders: Identification and care of the disorders. Journal of Cystic Fibrosis. 2024 Mar 19;

11. Smith AN, Barth ML, McDowell TL, Moulin DS, Nuthall HN, Hollingsworth MA, et al. A regulatory element in intron 1 of the cystic fibrosis transmembrane conductance regulator gene. J Biol Chem. 1996 Apr 26;271(17):9947–54.

12. Blotas C, Férec C, Moisan S. Tissue-Specific Regulation of CFTR Gene Expression. International Journal of Molecular Sciences. 2023 Jun;24(13):10678.

13. Corces MR, Trevino AE, Hamilton EG, Greenside PG, Sinnott-Armstrong NA, Vesuna S, et al. An improved ATAC-seq protocol reduces background and enables interrogation of frozen tissues. Nat Methods. 2017 Oct;14(10):959–62.

14. Tak YE, Horng JE, Perry NT, Schultz HT, Iyer S, Yao Q, et al. Augmenting and directing long-range CRISPR-mediated activation in human cells. Nat Methods. 2021 Sep;18(9):1075–81.

15. Skene PJ, Henikoff S. An efficient targeted nuclease strategy for high-resolution mapping of DNA binding sites. Reinberg D, editor. eLife. 2017 Jan 12;6:e21856.

16. Krijger PHL, Geeven G, Bianchi V, Hilvering CRE, de Laat W. 4C-seq from beginning to end: A detailed protocol for sample preparation and data analysis. Methods. 2020 Jan 1;170:17–32.

17. Geeven G, Teunissen H, de Laat W, de Wit E. peakC: a flexible, non-parametric peak calling package for 4C and Capture-C data. Nucleic Acids Research. 2018 Sep 6;46(15):e91.

18. Fulco CP, Nasser J, Jones TR, Munson G, Bergman DT, Subramanian V, et al. Activity-by-contact model of enhancer–promoter regulation from thousands of CRISPR perturbations. Nat Genet. 2019 Dec;51(12):1664–9.

19. Ibrahimi A, Velde GV, Reumers V, Toelen J, Thiry I, Vandeputte C, et al. Highly Efficient Multicistronic Lentiviral Vectors with Peptide 2A Sequences. Human Gene Therapy. 2009 Aug;20(8):845–60.

20. Mangeot PE, Risson V, Fusil F, Marnef A, Laurent E, Blin J, et al. Genome editing in primary cells and in vivo using viral-derived Nanoblades loaded with Cas9-sgRNA ribonucleoproteins. Nat Commun. 2019 Jan 3;10(1):45.

21. Banskota S, Raguram A, Suh S, Du SW, Davis JR, Choi EH, et al. Engineered virus-like particles for efficient in vivo delivery of therapeutic proteins. Cell. 2022 Jan 20;185(2):250–265.e16.

22. Nasser J, Bergman DT, Fulco CP, Guckelberger P, Doughty BR, Patwardhan TA, et al. Genome-wide enhancer maps link risk variants to disease genes. Nature. 2021 May;593(7858):238–43.

23. Moisan S, Berlivet S, Ka C, Le Gac G, Dostie J, Férec C. Analysis of long-range interactions in primary human cells identifies cooperative CFTR regulatory elements. Nucleic Acids Res. 2016 Apr 7;44(6):2564–76.

24. Nuthall HN, Moulin DS, Huxley C, Harris A. Analysis of DNase-I-hypersensitive sites at the 3’ end of the cystic fibrosis transmembrane conductance regulator gene (CFTR). Biochem J. 1999 Aug 1;341(Pt 3):601–11.

25. Blackledge NP, Carter EJ, Evans JR, Lawson V, Rowntree RK, Harris A. CTCF mediates insulator function at the CFTR locus. Biochemical Journal. 2007 Nov 14;408(2):267–75.

26. Simonis M, Klous P, Splinter E, Moshkin Y, Willemsen R, de Wit E, et al. Nuclear organization of active and inactive chromatin domains uncovered by chromosome conformation capture–on-chip (4C). Nat Genet. 2006 Nov;38(11):1348–54.

27. Rauluseviciute I, Riudavets-Puig R, Blanc-Mathieu R, Castro-Mondragon JA, Ferenc K, Kumar V, et al. JASPAR 2024: 20th anniversary of the open-access database of transcription factor binding profiles. Nucleic Acids Res. 2024 Jan 5;52(D1):D174–82.

28. Martinez-Ara M, Comoglio F, Steensel B van. Large-scale analysis of the integration of enhancer-enhancer signals by promoters. eLife [Internet]. 2024 Sep 10 [cited 2024 Oct 24];12. Available from: https://elifesciences.org/reviewed-preprints/91994

29. Osterwalder M, Barozzi I, Tissières V, Fukuda-Yuzawa Y, Mannion BJ, Afzal SY, et al. Enhancer redundancy provides phenotypic robustness in mammalian development. Nature. 2018 Feb;554(7691):239–43.

30. Kim S, Wysocka J. Deciphering the multi-scale, quantitative *cis-*regulatory code. Molecular Cell. 2023 Feb 2;83(3):373–92.

31. van Mierlo G, Pushkarev O, Kribelbauer JF, Deplancke B. Chromatin modules and their implication in genomic organization and gene regulation. Trends in Genetics. 2023 Feb 1;39(2):140–53.

32. McCarthy VA, Ott CJ, Phylactides M, Harris A. Interaction of intestinal and pancreatic transcription factors in the regulation of CFTR gene expression. Biochim Biophys Acta. 2009;1789(11–12):709–18.

33. Smith EM, Lajoie BR, Jain G, Dekker J. Invariant TAD Boundaries Constrain Cell-Type-Specific Looping Interactions between Promoters and Distal Elements around the CFTR Locus. Am J Hum Genet. 2016 Jan 7;98(1):185–201.

34. Zhang Z, Ott CJ, Lewandowska MA, Leir SH, Harris A. Molecular mechanisms controlling CFTR gene expression in the airway. J Cell Mol Med. 2012 Jun;16(6):1321–30.

35. Collobert M, Bocher O, Le Nabec A, Génin E, Férec C, Moisan S. CFTR Cooperative Cis-Regulatory Elements in Intestinal Cells. Int J Mol Sci. 2021 Mar 5;22(5):2599.

36. Hung TC, Kingsley DM, Boettiger AN. Boundary stacking interactions enable cross-TAD enhancer–promoter communication during limb development. Nat Genet. 2024 Feb;56(2):306–14.

37. Yang JH, Hansen AS. Enhancer selectivity in space and time: from enhancer–promoter interactions to promoter activation. Nat Rev Mol Cell Biol. 2024 Feb 27;1–18.

38. Mummey HM, Elison W, Korgaonkar K, Elgamal RM, Kudtarkar P, Griffin E, et al. Single cell multiome profiling of pancreatic islets reveals physiological changes in cell type-specific regulation associated with diabetes risk. bioRxiv. 2024 Aug 6;

39. Hecker D, Behjati Ardakani F, Karollus A, Gagneur J, Schulz MH. The adapted Activity-By-Contact model for enhancer–gene assignment and its application to single-cell data. Bioinformatics. 2023 Feb 1;39(2).

40. Hoellinger T, Mestre C, Aschard H, Le Goff W, Foissac S, Faraut T, et al. Enhancer/gene relationships: Need for more reliable genome-wide reference sets. Front Bioinform. 2023 Feb 24;3.

41. Ying P, Chen C, Lu Z, Chen S, Zhang M, Cai Y, et al. Genome-wide enhancer-gene regulatory maps link causal variants to target genes underlying human cancer risk. Nat Commun. 2023 Sep 25;14(1):5958.

42. Malfait J, Wan J, Spicuglia S. Epromoters are new players in the regulatory landscape with potential pleiotropic roles. Bioessays. 2023 Oct;45(10):e2300012.

43. Smith AN, Wardle CJ, Harris A. Characterization of DNASE I hypersensitive sites in the 120kb 5’ to the CFTR gene. Biochem Biophys Res Commun. 1995 Jun 6;211(1):274–81.

44. Xie F, Armand EJ, Yao Z, Liu H, Bartlett A, Behrens MM, et al. Robust enhancer-gene regulation identified by single-cell transcriptomes and epigenomes. Cell Genomics. 2023 Jul 12;3(7):100342.

45. Merkenschlager M, Nora EP. CTCF and Cohesin in Genome Folding and Transcriptional Gene Regulation. Annu Rev Genom Hum Genet. 2016 Aug 31;17(1):17–43.

46. Brosh R, Laurent JM, Ordoñez R, Huang E, Hogan MS, Hitchcock AM, et al. A versatile platform for locus-scale genome rewriting and verification. Proceedings of the National Academy of Sciences. 2021 Mar 9;118(10):e2023952118.

47. Xu Y, Nipper MH, Dominguez AA, Ye Z, Akanuma N, Lopez K, et al. Reconstitution of human PDAC using primary cells reveals oncogenic transcriptomic features at tumor onset. Nat Commun. 2024 Jan 27;15(1):818.

