## Supplementary data for "New insights into the *cis*-regulation of the *CFTR* gene in pancreatic cells"

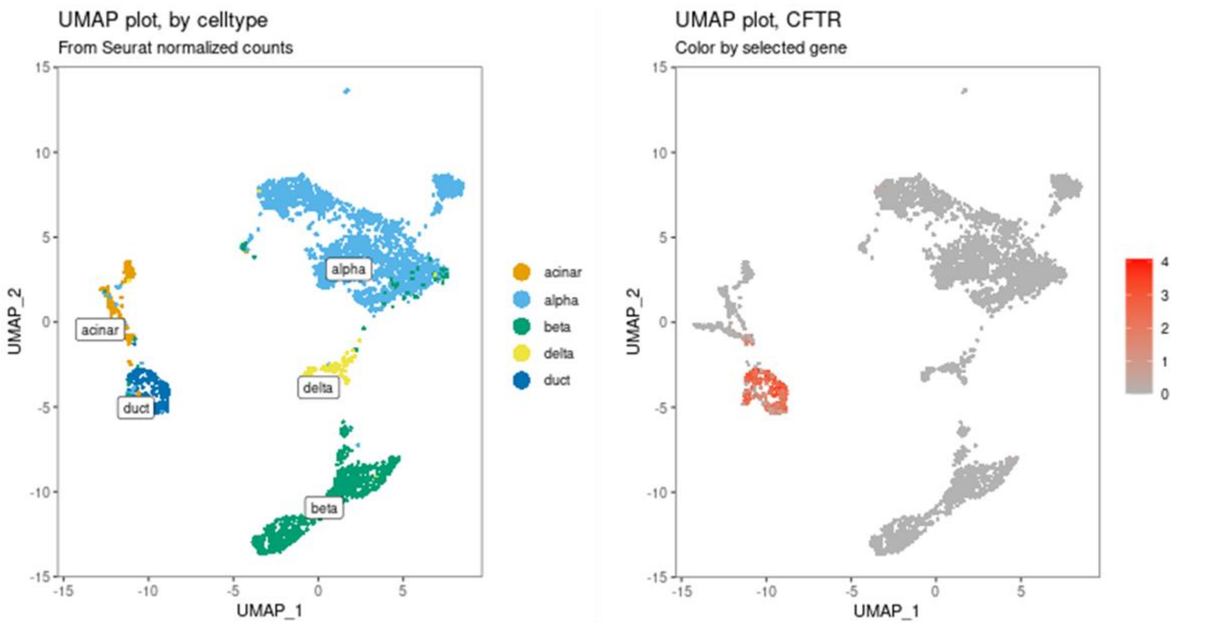


***Figure S1. Single-cells CFTR expression.***

*Single-cell RNA-seq data from the PancrESS dataset [40]. Uniform Manifold Approximation and Projection (UMAP) of transcript abundances. On the left: cells are color-coded by cell type, demonstrating clustering by cell type. On the right: cells expressing the gene of interest, CFTR, are highlighted in red, with the color intensity representing transcript abundance levels.*

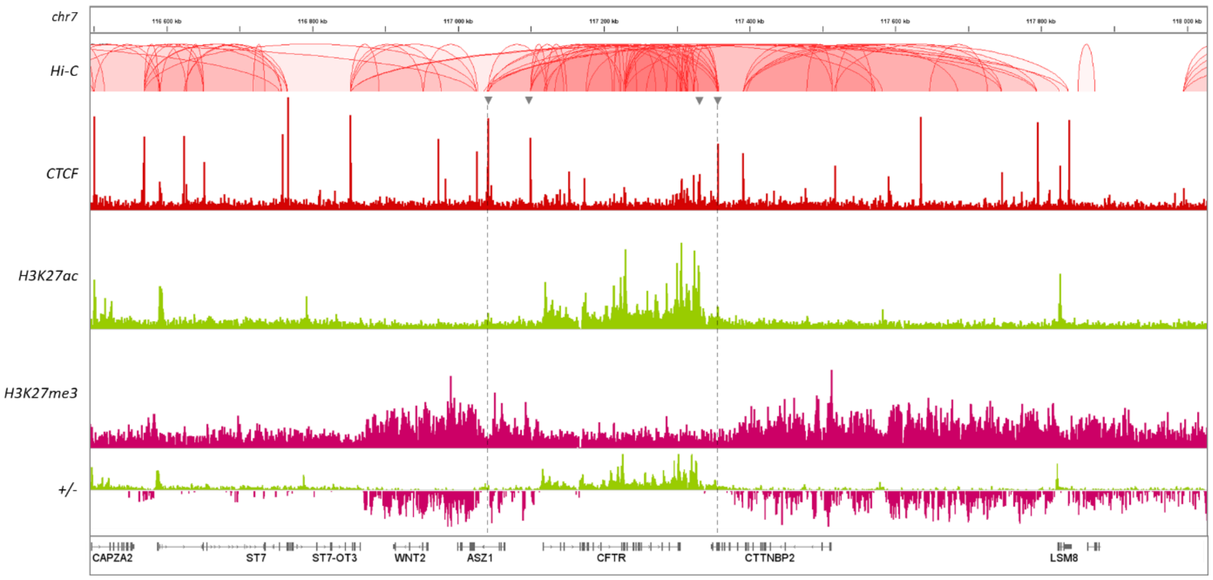


***Figure S2. Chromatin data from Caco-2 cells.***

*Hi-C data provide information of genome-wide chromatin interaction. CTCF immunoprecipitation allows to map CTCF binding. H3K27ac and H3K27me3 provide information of the chromatin state, acetylation is for active chromatin and trimethylation is for repress chromatin. TAD boundaries and enhancer blocking sites are indicated by grey arrow. We noticed the correlation of chromatin state and epigenetic marks.*

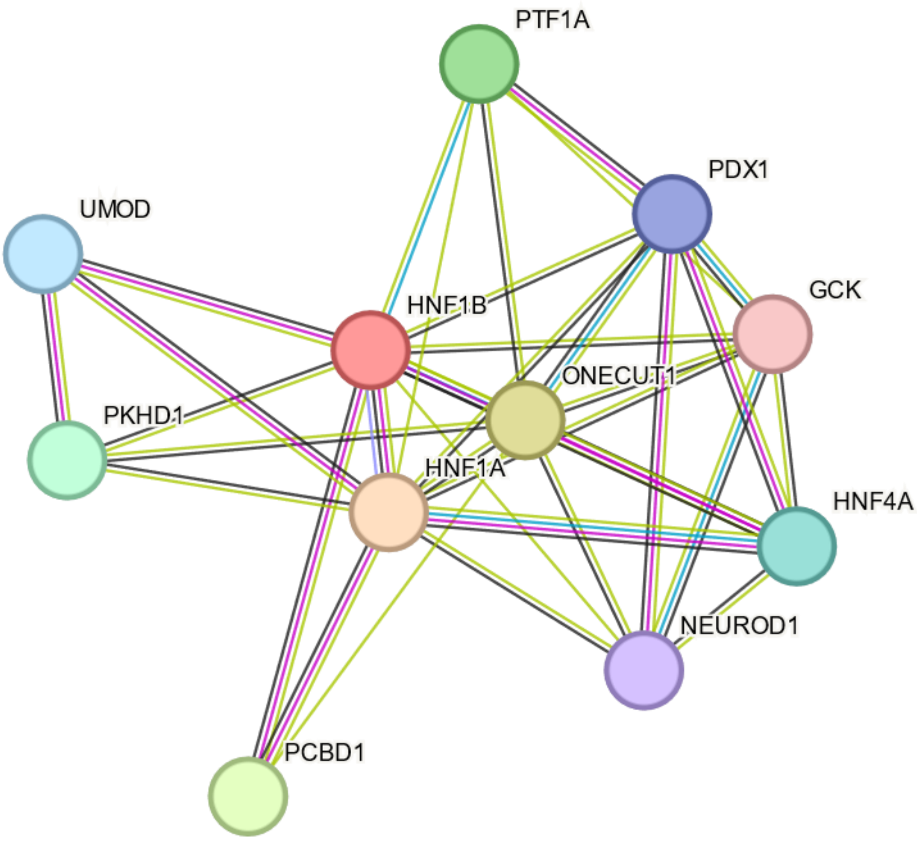


***Figure S3. Transcription Factor Interaction Network.***

*The STRING database provides an interaction network with HNF1b as the bait. The edges represent protein-protein associations.*

***Table S1. PeakC analysis of 4C data from Capan-1 cells with the CFTR promoter as bait (chr7:117,039,879-117,356,81, hg19)***

| | CRE | Genomic coordinates (hg19) |
|---|---|---|
| **1** | -80.1 kb | chr7:117042598-117048074 |
| **2** | - | chr7:117057283-117064712 |
| **3** | Promoter | chr7:117113481-117125179 |
| **4** | +15.6 kb | chr7:117310255-117329893 |

***Table S2. 4C-seq primers***

| Primers Name | Primers Sequence |
|---|---|
| **4C_pCFTR_reading** | TACACGACGCTCTTCCGATCTGTCTTTCCATCATGAAATGCAC |
| **4C_pCFTR_nonreading** | ACTGGAGTTCAGACGTGTGCTCTTCCGATCTGAATTAAGCTCCAAAGAGGATC |

*Table S3. Primers for plasmid constructs*

| CRE | Forward/reverse | Primers sequence |
|---|---|---|
| -44 KB | F | AAATCGATAAGGATCCGGTCATAGATGTGGAACATTCTG |
| | R | GATCAGATGTGGGATCCCCGCAAAATCTGCCATCATC |
| -35 KB | F | AAATCGATAAGGATCCCTCATGTAGCAGGCATTTGTG |
| | R | ATCGGTCGACGGATCCGTGTTTCCAATTTCTACCCAGAC |
| +15,6 KB | F | AAATCGATAAGGATCCCAGTGCCTGGCCAAGAATC |
| | F | ATCGGTCGACGGATCCGAGGTATCAGAAATCAGTGTGAG |
| +37,7 KB | R | AAATCGATAAGGATCCTACCACGTAAGTGAGATTATGTGG |
| | F | ATCGGTCGACGGATCCGAGTGGCTTCAAACCTTACTTATTC |
| -44 KB / -35 KB | R | CTAGCCCGGGCTCGAGGGTCATAGATGTGGAACATTCTG |
| | F | CAAATGCCTGCTACATGAGCCGCAAAATCTGCCATCATC |
| | F | GATGATGGCAGATTTTGCGGCTCATGTAGCAGGCATTTG |
| | R | GATCGCAGATCTCGAGCCAGAAGTTCACAAGACTG |
| +15,6 KB / +37,7 KB | F | AAATCGATAAGGATCCCAGTGCCTGGCCAAGAATC |
| | R | CATAATCTCACTTACGTGGTAGGAGGTATCAGAAATCAGTGTG |
| | F | CACACTGATTTCTGATACCTCCTACCACGTAAGTGAGATTATG |
| | R | AAATCGATAAGGATCCTACCACGTAAGTGAGATTATGTGG |
| -3,4 KB | F | AAATCGATAAGGATCCCTTTAGCACCCAAGCTCTTG |
| | R | ATCGGTCGACGGATCCCCTTATGGTTGTTGGTCACTTAC |
| Intron 11 | F | AAATCGATAAGGATCCCTTAGTTCTCATGCTTTCTAGTGG |
| | R | ATCGGTCGACGGATCCGCAGTGGAGGATGGTTTGAAAG |
| Intron 12 | F | AAATCGATAAGGATCCTGGAGAAGGTGGAATCACACTG |
| | R | ATCGGTCGACGGATCCGAAGACAGTATGCAAGAGCTACAT |
| Intron 18 | F | AAATCGATAAGGATCCCTCCCCATTTCCCTCTCTCC |
| | R | ATCGGTCGACGGATCCGCTGGGCATTCGTCTTGGAG |
| Intron 24 | F | AAATCGATAAGGATCCGCATTTCTCACTCTGGCTGG |
| | R | ATCGGTCGACGGATCCCCTGTCCAAACTAAGAAGACTTC |
| Intron 12-24 | F | AAATCGATAAGGATCCTGGAGAAGGTGGAATCACACTG |
| | R | TACTGTCTTCGGATCCTCGACGCATTTCTCACTCTGGCTGG |
| | F | AAGGGCATCGTCGACCCTGTCCAAACTAAGAAGACTTC |
| | R | ATCGGTCGACGGATCCCCTGTCCAAACTAAGAAGACTTC |
| Intron 26 | F | AAATCGATAAGGATCCATCTGAGCCATGTGGTGAGGTTGA |
| | R | ATCGGTCGACGGATCCTTCCTTTGAGCCTGTGCCAGTTTC |
| +484,2 KB | F | AAATCGATAAGGATCCGTACGAAGTATCAACTAGGAGAGC |
| | R | ATCGGTCGACGGATCCTCTTAGATAATTTAAAAAGCC |
| +507,6 KB | F | AAATCGATAAGGATCCGTGCCACAGAAAAAGATGGCC |
| | R | ATCGGTCGACGGATCCGAATAATGTAAGAAGGTATATAG |
| +513,7 KB | F | AAATCGATAAGGATCCGAGCATTACTTTGAGAGGTTTCGAC |
| | R | ATCGGTCGACGGATCCGTGTCTTCCGCTAATGAAAGAAC |

*Table S4. gRNA and PCR primers for CRISPR/cas9 deletion of -44 kb region*

| **gRNA name** | **gRNA sequence** |
|---|---|
| **sgRNA-CRE1_F** | CAGAGAGGCCTGGGCTATCC |
| **sgRNA-CRE1_R** | GATATCAGACAACAAGTCTA |
| **PCR primers name** | **Primers sequence** |
| **Cis_CFTR_-44kb_Ext_F** | GGAAACAGGTAAGTTTTGTTCTG |
| **Cis_CFTR_-44kb_Ext_R** | GCTTTGTTCTCTGACCTAACC |
| **qPCR primers name** | **Primers sequence** |
| **CFTR_qPCR_F** | ATGCCCTTCGGCGATGTTTT |
| **CFTR_qPCR_R** | TGATTCTTCCCAGTAAGAGAGGC |
| **Actin_qPCR_F** | CTGGAACGGTGAAGGTGACA |
| **Actin_qPCR_R** | AAGGGACTTCCTGTAACAATGCA |
